# Remodeling of the secretory pathway is coordinated with *de novo* membrane formation in budding yeast gametogenesis

**DOI:** 10.1101/2023.07.10.548399

**Authors:** Yasuyuki Suda, Hiroyuki Tachikawa, Tomomi Suda, Kazuo Kurokawa, Akihiko Nakano, Kenji Irie

## Abstract

Gametogenesis in budding yeast involves a large-scale rearrangement of membrane traffic to allow *de novo* formation of a membrane, called the prospore membrane (PSM). However, the mechanism underlying this event is not fully elucidated. Here, we show that the number of endoplasmic reticulum exit sites (ERES) per cell fluctuates and switches from decreasing to increasing upon the onset of PSM formation. Reduction in ERES number, presumably accompanying a transient stall in membrane traffic, resulting in the loss of preexisting Golgi apparatus from the cell, was followed by local ERES regeneration, leading to Golgi reassembly in nascent spores. We have revealed that protein phosphatase-1 (PP-1) and its development-specific subunit, Gip1, promote ERES regeneration through Sec16 foci formation. Furthermore, *sed4*Δ, a mutant with impaired ERES formation, showed defects in PSM growth and spore formation. Thus, ERES regeneration in nascent spores facilitates the segregation of membrane traffic organelles, leading to PSM growth.

## Introduction

Gametogenesis of budding yeast is a developmental process consisting of meiosis and spore formation, in which chromosomes stored within the nucleus undergo a single round of replication followed by two successive rounds of segregation (meiosis I and II) to produce haploid gametes. In meiosis II, a newly generated membrane, the prospore membrane (PSM), is formed at the cytoplasmic surface of the spindle pole body (SPB) embedded in the nuclear membrane, and the haploid set of chromosomes in the nucleus is engulfed by the growing PSM, along with organelles and cytosol^1^. This process is triggered by starvation, accompanied by a massive remodeling in the mother cell, is carried out through the strict control of transcription and translation, and depends on membrane traffic^1,2^.

In mitotic cell growth, both secretory cargo molecules and lipids synthesized in the endoplasmic reticulum (ER) are first transported to the Golgi apparatus and then sorted to their final destinations, post-Golgi compartments and the extracellular space. The itinerary of this default secretory pathway is dramatically altered during meiosis, and its destination is changed to the PSM^3^. In this case, the formation of the PSM is initiated by converting the molecular mechanisms of post-Golgi vesicle fusion with the plasma membrane in mitotic cells into those at cytoplasmic surface of the SPB^4^. The mother cell requires a large supply of lipids to generate the large membrane structures from scratch that later become the gamete’s plasma membrane. To accomplish this, not only the conversion of membrane traffic but also lipid transport by Vps13, a newly identified lipid transporter, and its adaptor complex is required^5–9^. The Vps13 complex is assumed to transport lipids directly from the ER to the PSM through its hydrophobic tunnel-like structure^5,10^. Although various studies have shed light on the mechanisms of nascent PSM formation, the integrated regulation of membrane traffic throughout the cell during meiotic progression remains unclear.

Transport of cargo proteins from the ER is mediated by the COPII transport carrier that is formed at specialized ER subdomains called ER exit sites (ERES)^11^. The COPII carrier consists of inner coat Sec23-Sec24 and outer coat Sec13-Sec31 complexes, which are recruited to the membrane by the active Sar1-GTPase^12^. ERES are preferentially formed at the high-curvature domain of the ER, which is composed of fenestrated sheets and tubules^13^. Sec16 is a large peripheral membrane protein and a key molecule for ERES organization, since its dysfunction causes failure of COPII-mediated transport as well as ERES formation^14,15^. The potential function of Sec16 in ERES organization is widely conserved throughout eukaryotes, but its regulation differs among organisms^16^. In mammals, during mitosis, the ERES collapse and the Golgi fragment disperses into the cytoplasm, thereby the secretory pathway is transiently attenuated^17,18^. In *Drosophila melanogaster*, upon starvation, Sec16 is released from the ER membrane and sequestered in a phase-separated granule with several COPII-forming factors^19^. This sequestration is bidirectional and is assumed to function as a reservoir to initiate COPII formation quickly after recovery from starvation. Mammalian cell division and the starvation response of flies are processes that involve large-scale remodeling, in which cells employ integrated regulation of membrane traffic via its starting point, the ERES. Gametogenesis in budding yeast also involves cellular remodeling, but it is not well understood whether ERES formation is similarly regulated.

We previously showed that the cortical ER, a subset of the ER closely associated with the plasma membrane, is reorganized upon PSM formation and that the PSM acts as a border to keep the Golgi apparatus inside^20^. More recently, the molecular mechanisms responsible for the morphology of the ER during meiosis and for the distribution of the mitochondria have also been elucidated^21,22^. Although these organellar remodeling processes are shown to be a system to exclude unnecessary organelles rather than an active transport machinery toward the gametes, they are crucial to the cell, as it is inherently linked to cell rejuvenation by eliminating age-related factors from the gametes^2,23^. In this case, the PSM provides a barrier between age-related factors such as damaged organelles and essential organelles including nucleus to be inherited by the next generation. Thus the progeny become protected by the robust spore wall formed in the lumen of PSMs, while the remaining age-related factors are degraded by autophagy and subsequent vacuolar rupture^24^.

In this study, we revealed the dynamics of ERES during gametogenesis and its involvement in PSM formation; ERES formation is transiently inactivated during meiosis, which causes a transient pause in membrane traffic leading to the Golgi disassembly. Subsequently, ERES regeneration is locally activated in the region surrounded by the PSM in meiosis II, facilitating re-assembly of the Golgi and PSM growth. Analysis of mutants defective in PSM formation revealed that Gip1, a meiosis-specific targeting subunit of PP1^25,26^, functions in ERES remodeling. Its direct contribution to PSM formation was previously unknown. PP1-dependent ERES remodeling occurs through Sec16. The analysis of factors involved in COPII-carrier formation suggests that the reorganization of membrane traffic in presumptive spores is the system responsible for the segregation of membrane traffic into gametes, and that lipid supply through the secretory pathway contributes to PSM growth. Overall, our study indicates that developmental regulation of membrane traffic coordinates *de novo* membrane formation and organellar segregation to the progeny.

## Results

### ERES numbers fluctuate in meiosis

Meiotic development includes faithful inheritance of essential organelles such as the nucleus, mitochondria, and ER for generation of spore gametes. To characterize morphologies of the organelles for membrane traffic upon meiotic development in detail, time-course observation of the ER was performed in cells expressing the transmembrane domain of ER-localized tethering protein, Scs2, fused with mNeonGreen (mNG-Scs2TM) and chromatin (Htb2-mCherry) (Fig. 1A). In premeiotic cells, reticular cortical ER expanded underneath the plasma membrane and was retained as in vegetatively growing cells (Fig. 1A. 0-3 h). Detachment of cortical ER from the plasma membrane and expansion of cytoplasmic ER were observed before the first meiotic division (Fig. 1A. 4 h). Then cortical ER were lost as meiosis progressed, presumably being absorbed into the nuclear ER (Fig. 1A. 5 h-). In metaphase II to anaphase II cells, ER signals were detected in association with dividing nuclei (Fig. 1A. 6-9 h), similar to the previous observation^20^. Subsequently, regeneration of the cortical ER underneath the spore plasma membrane was seen during completion of spore maturation (Fig. 1A. 10 h), consistent with the prior data^21^.

**Figure 1.**
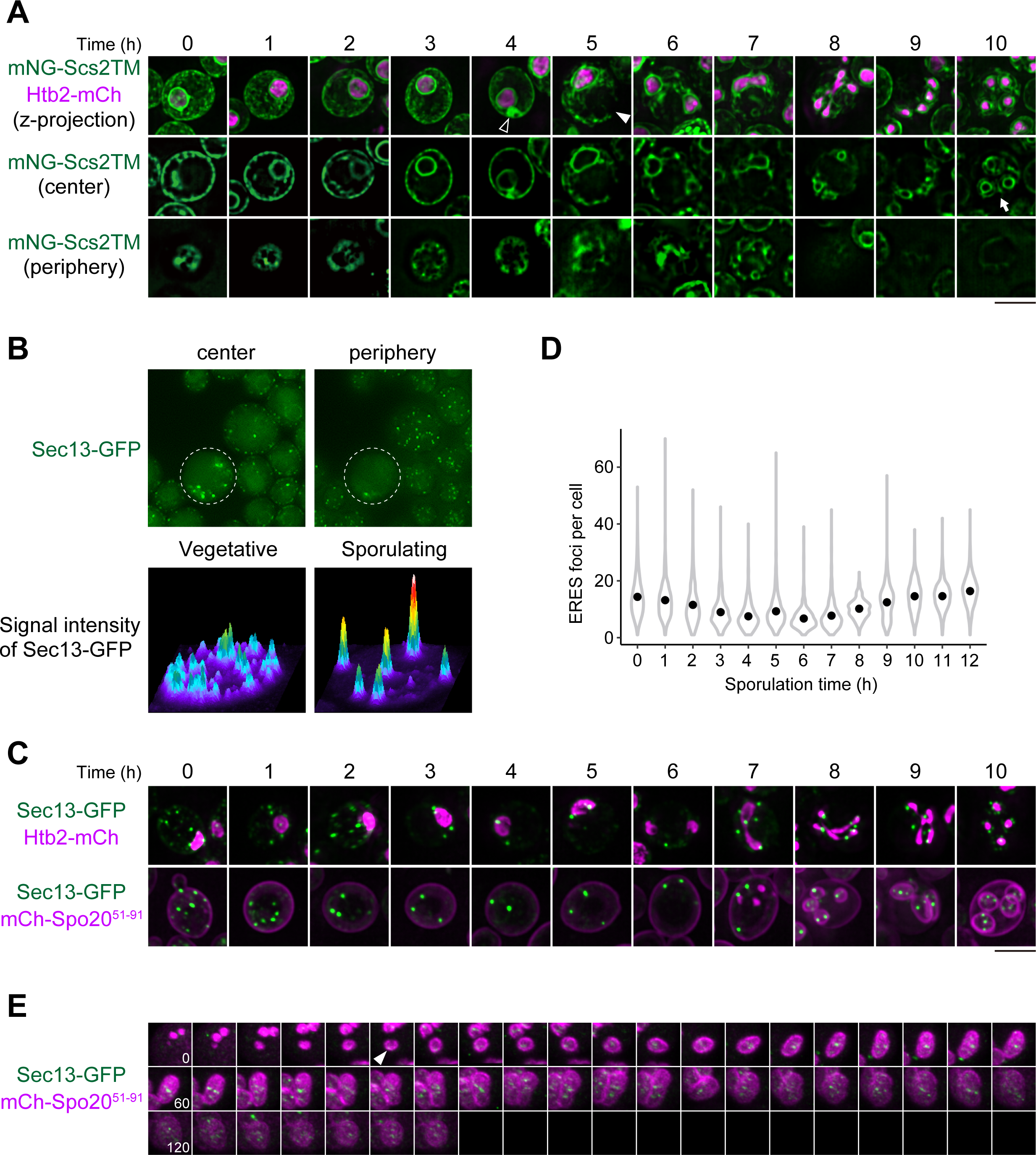
Distribution of ERES during meiotic progression. (A) Maximum intensity projections (top) or single planes at the center (middle) or periphery (bottom) of cells expressing mNG-Scs2TM (ER marker) and Htb2-mCherry (nucleoplasm marker) at each time point in meiosis. Detachment of cortical ER (closed arrowhead), expansion of cytoplasmic ER (open arrowhead), and regeneration of cortical ER within the spores (arrow) are shown. (B) Single planes at the periphery (upper left) or center (upper right) of sporulating (dashed line) and vegetative (others) cells expressing Sec13-GFP (ERES marker). Signal intensity profiles for Sec13-GFP in vegetative and sporulating cells are also shown (bottom). (C) Maximum intensity projections of cells expressing Sec13-GFP with either Htb2-mCherry or mCherry-Spo20^51–91^ (PSM marker) at each time point after induction of meiosis. (D) Quantification of the experiments as in upper images of panel C. Number of Sec13-GFP dots were counted (n > 300 cells /h). Data are presented as violin plots with median. (E) Regeneration of ERES in presumptive spores. Time-lapse images of single presumptive spores from cells expressing Sec13-GFP and mCherry-Spo20^51–91^ in meiosis. Kymograph from Movie S1 is shown.

ERES are sites for the assembly of COPII transport carriers and are formed preferentially at the high-curvature domain of the ER^13^. Observation of ERES in pre-meiotic cells, visualized by Sec13-GFP, showed numerous spots in the peripheral region of the cell. In contrast, as meiosis progresses, ERES disappeared from cell periphery and scattered in the cytoplasm (Fig. 1B). To analyze ERES dynamics in meiotic development, time-course observation of ERES was also performed with chromatin marked by Htb2-mCherry (Fig. 1C). Numerous punctate structures of ERES were observed in premeiotic cells like mitotic cells, but their number gradually decreased and then increased after metaphase II (Fig. 1D). Although the number of ERES fluctuated during meiotic progression, the average signal intensities of Sec13-GFP were relatively stable (Fig. S1A, S1B). We reasoned that the observed change in the number of ERES could correspond to the activity of COPII generation. In the fruit fly, *D. melanogaster*, amino acid starvation leads to the coalescence of Sec16, the scaffold protein for the assembly of COPII transport carriers, into large membrane-less organelles called sec bodies, which are thought to act as a reservoir for ERES in harsh conditions^19^. Under the condition for budding yeast meiosis, Sec16-GFP was similarly distributed and colocalized with Sec13-mCherry (Fig. S1C, S1D). By analyzing Sec13-GFP dynamics by fluorescence recovery after photobleaching (FRAP), we showed that Sec13-GFP molecules exchange rapidly between ERES and the cytoplasm in vegetatively growing cells. The half-time to recovery was 4.6 s (+/- 2.4 s). By contrast, in cells undergoing meiosis, the half-time to recovery for meiosis I and II was 18.8 s (+/- 16.9 s) and 26.5 s (+/- 21.6 s), respectively (Fig. S1E and S1F). These data indicate that the activity of ERES persists, but its dynamics significantly slowed down in the progression of meiosis.

### ERES forms *de novo* in the spore cytoplasm

In anaphase II cells, ERES always localized in association with dividing chromosomes and were precisely segregated into presumptive spores (Fig. 1C). We next investigated the proper timing of ERES segregation into nascent spores. The PSM is formed by the coalescence of post-Golgi vesicles at the cytoplasmic face of SPBs^1^. Based on time-course observation of PSM perimeters, PSM growth was categorized into the initial fusion process, horseshoe, elongation, and mature stages^8^. ERES segregation was found in cells with the horseshoe stage (Fig. 1C). To further characterize the segregation of ERES during PSM formation, we performed 3D live-cell imaging of ERES and PSM dynamics by super-resolution confocal live imaging microscopy (SCLIM)^27,28^. ERES first appeared inside the PSM at the stage (∼12 min) in which the PSM showed horseshoe-like morphology, then ERES proliferated inside PSMs during the elongation stage until the completion of PSM maturation (Fig. 1E and Movie S1). Taken together, these data reveal that the collapse of ERES leads to the decrease of their number in the cell during meiotic progression until the onset of PSM formation, then they are formed *de novo* and proliferate in the cytoplasm of nascent spores.

### ERES dynamics coincide with the Golgi disappearance and reappearance

In yeast, the Golgi can form *de novo*, as COPII vesicles arising from ERES fuse to form a post-ER compartment and then presumably mature into ERGIC (ER-Golgi intermediate compartment), then *cis*-cisterna of the Golgi^11,29,30^. We have previously shown that the nascent PSM functions as a diffusion barrier for the Golgi^20^, however detailed dynamics of the ER-Golgi unit during meiosis have not been examined. To this end, we investigated whether the collapse and regeneration of the Golgi in cells are also observed during meiotic progression and PSM formation. To observe the Golgi apparatus, we chose the fluorescently tagged Golgi-resident proteins, Grh1-2xyEGFP, a homolog of the mammalian Golgi reassembly stacking protein 65 kD (GRASP65) that can be used for ERGIC marker, and Mnn9-sfGFP, a *cis*-Golgi resident mannosyltransferase^30–32^. Although Grh1-2xyEGFP dots were seen in PSMs in most of the cells irrespective of the stages of the PSM growth, Mnn9-sfGFP dots were observed only in the cytoplasmic space of elongation stage of PSMs, but not in that of horseshoe-stage of PSMs (Fig. 2A). Note that Mnn9-sfGFP signals showed ER-like pattern in the cells with horseshoe-stage of PSMs (Fig 2A, arrowhead, Fig S2A). These observations could reflect both disassembly and regeneration of the Golgi during the formation of PSMs, which might be caused on the change of ERES number in meiosis. We have recently observed that, upon secretion block such as by the brefeldin A (BFA) treatment, ERGIC components including Grh1 signal accumulated as clusters and most of Golgi-resident proteins including glycosyltransferases were absorbed into the ER^30^, which is exactly like the case with this observation during meiosis II. Furthermore, similar distributions of Grh1-2xyEGFP and Mnn9-sfGFP signals were also observed at the restrictive temperature in *sec16-2* mutant cells (Fig. 2B), in which secretion was blocked between the ER and the Golgi^33^. These findings suggest that temporal secretion block also occur due to the decrease in ERES number during meiotic progression, and the Golgi reassemble within the spore cytoplasm presumably from the fusion of COPII vesicles arising from ERES.

**Figure 2.**
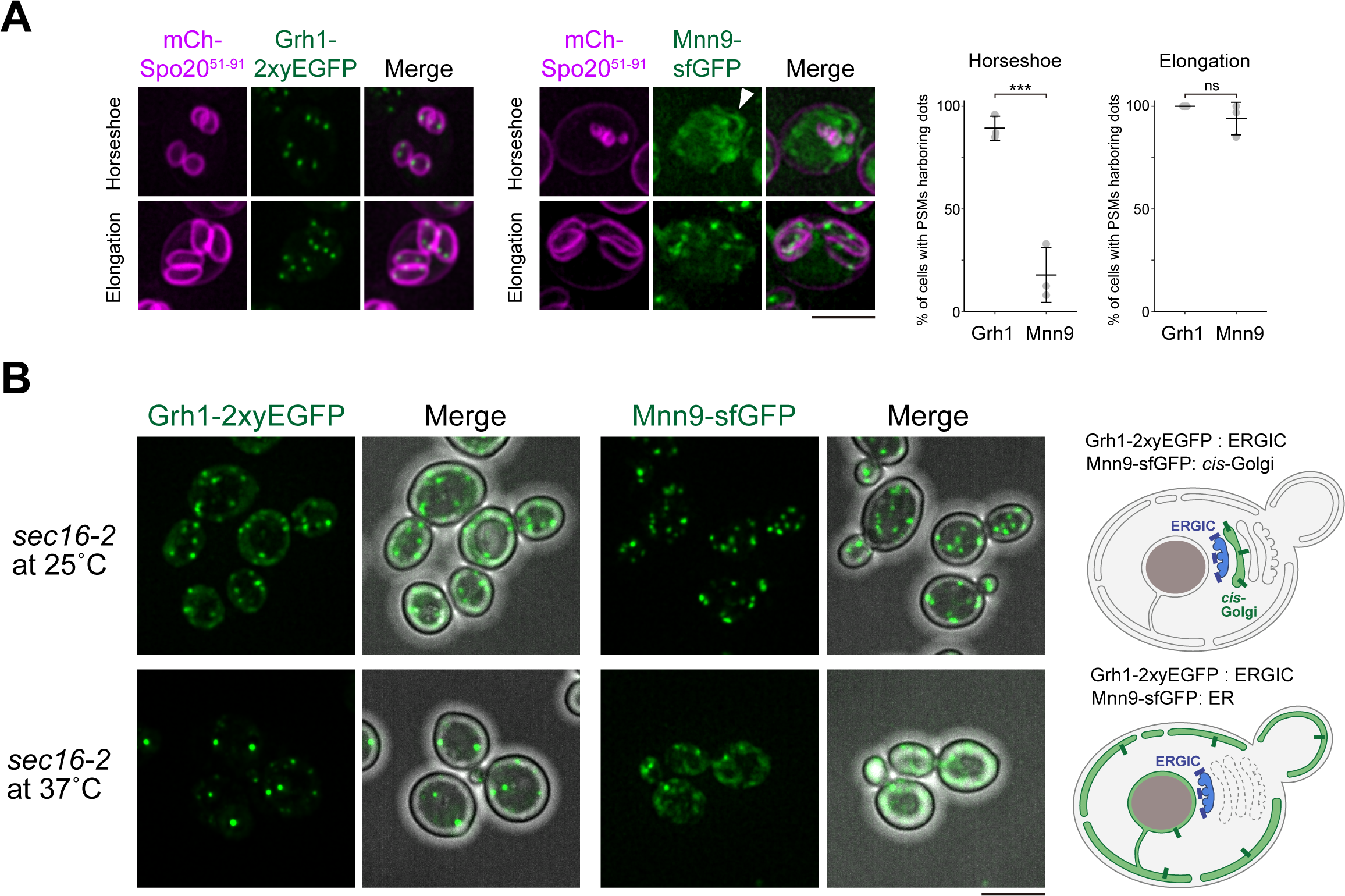
Regeneration of the Golgi in presumptive spores. (A) Maximum intensity projections of the cells expressing ERGIC marker Grh1-2xyEGFP and *cis*-Golgi localizing enzyme Mnn9-sfGFP with PSM marker mCherry-Spo20^51–91^ at each stage of PSM progression. Graphs showing the percentage of cells with PSMs harboring indicated markers at horseshoe and elongation stages of PSM growth. More than 100 cells were analyzed in three independent trials. *P* values were calculated with unpaired two-tailed Welch’s *t*-test. ***, *P* < 0.01, ns, not significant. (B) Maximum intensity projections of *sec16-2* mutant cells expressing Grh1-2xyEGFP or Mnn9-sfGFP at permissive (25°C) or restrictive (37°C) temperature. Merge panels are shown with bright field image. Schematics of Grh1-2xyEGFP and Mnn9-sfGFP distribution at indicated conditions are shown on the right. The Golgi cisternae are shown in a stack for simplicity.

### Determinants of ER shape are important for change in ERES numbers

ERES formation is preferred at the high-curvature region of the ER^13^. As reticulons and DP1/Yop1 are factors stabilizing the network-like morphology of the ER^34^, ERES accumulate at the edge of the expanded sheet of the ER in *rtn1*Δ *rtn2*Δ *yop1*Δ cells^13^. In meiosis, reticulon-mediated ER curvature has been reported to lead directly or indirectly to the cortical detachment and cabling of the ER^21^. To test whether the changes in ERES numbers are also affected in this situation, we analyzed the morphology of the ER and the distributions of ERES numbers in *rtn1*Δ *rtn2*Δ *yop1*Δ cells. Consistent with the previous observation^21^, there was no expansion of the cytoplasmic ER or absorption into the nuclear ER, and rather smooth, sheet-like structures were retained near the cortical region of the cell (Fig. 3A, 3B). Thus, the cytoplasmic expansion is the result of the detachment of cortical ER. The number of ERES was not decreased but rather persisted in metaphase II (Fig. 3C), corresponding to the observed defect of the ER morphology in *rtn1*Δ *rtn2*Δ *yop1*Δ cells. These results suggest that the number and distribution of ERES are linked, at least in part, to the morphological change of the ER in meiotic development.

**Figure 3.**
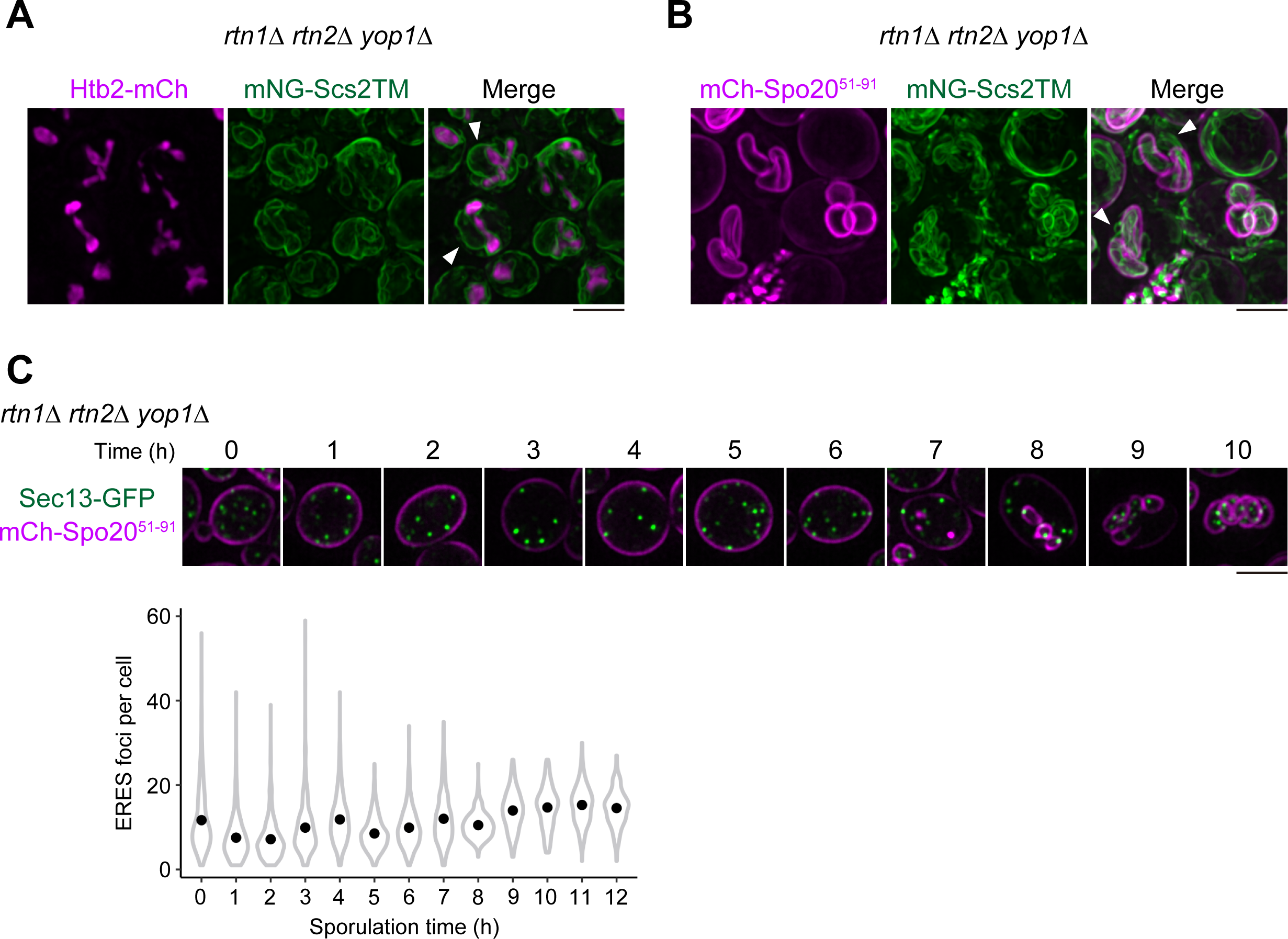
ER-shaping determinants are important for the distribution of ERES numbers in meiosis. (A-B) Maximum intensity projections of anaphase II *rtn1*Δ *rtn2*Δ *yop1*Δ cells expressing (A) mNG-Scs2TM and Htb2-mCherry or (B) mNG-Scs2TM and mCherry-Spo20^51–91^. (C) Maximum intensity projections of *rtn1*Δ *rtn2*Δ *yop1*Δ cells expressing Sec13-GFP with mCherry-Spo20^51–91^ at each time point after induction of meiosis. Lower graph shows quantification of the experiments in upper images. Number of Sec13-GFP dots were counted (n > 300 cells /h). Data are presented as violin plots with median.

### ERES regeneration within spores requires a meiosis-specific PP1 subunit

To clarify the molecular basis for ERES regeneration within the spore cytoplasm, we carried out visual inspection among the mutants displaying defects in the morphological development of the PSM, including *gip1*Δ, *vps13*Δ, *spo71*Δ, *spo73*Δ, *sma2*Δ, *spo1*Δ, and *spo19*^8,26,35–37^ ERES foci were observed inside the PSM (Fig. 4A, upper panels, and 4B). Gip1 is a sporulation-specific targeting subunit for Glc7, the sole catalytic subunit of PP1 in budding yeast, and *gip1*Δ showed multiple defects in spore development such as septin organization, PSM growth, and spore wall formation^25,26^. PSM extension is facilitated independently by unknown functions of PP1-Gip1 and Vps13-mediated lipid transport^25^. Although a similar PSM growth defect has been shown in mutants for Vps13 and its adaptor complex Spo71-Spo73^8^, ERES distributed inside the PSM in these mutant cells (Fig. 4A, lower panels, and 4B). Time-course analyses of the numbers of ERES in *gip1*Δ and *vps13*Δ mutants were indistinguishable from that in wild-type cells (Fig. 4C and Fig. 1C, 1D). In *gip1*Δ cells, the initial process of PSM formation was normal, but the elongation was defective, stopping at the horseshoe-stage (PSM perimeter of almost 6 µm) (Fig. 4D, *gip1*Δ). Whereas ERES regeneration was observed in most of the wild-type cells with a PSM perimeter of at least 2 µm (Fig. 4E, wt), the *gip1*Δ PSM failed to contain ERES, even in cells with PSM lengths greater than 2 µm perimeters (Fig. 4E, *gip1*Δ). To assess whether ERES regeneration occurs in *gip1*Δ, we conducted 3D live-cell imaging using SCLIM. Observation of Sec13-GFP in *gip1*Δ cells showed no ERES signals accumulate within the spore cytoplasm along with the defect in PSM elongation (Fig. 5A and Movie S2). Quantification of the signal intensity ratio of Sec13-GFP between inside the PSM and the entire cell, clearly showed the *de novo* formation of ERES within the PSM in wild-type cells but not in *gip1*Δ cells (Fig. 5B). No characteristic defect in the ER morphology was found in the *gip1*Δ mutant during meiosis (Fig. 5C). In *gip1*Δ cells, regeneration of the Golgi was completely abolished in all PSMs formed (Fig. 5D). These results indicate that the temporal secretion block is not resumed in *gip1*Δ cells. To confirm this, delivery of the cargo, sporulation-specific subtilisin-like protease Osw3^38^, to the PSM was assessed. Osw3-Envy reached PSMs in wild-type cells, however Osw3-Envy signals were observed predominantly at the ER in *gip1*Δ cells (Fig. 5E), suggesting that the secretion did not resume in *gip1*Δ cells. Collectively, we propose that Gip1 is required for ERES regeneration in nascent spores to reestablish membrane traffic, leading to PSM elongation.

**Figure 4.**
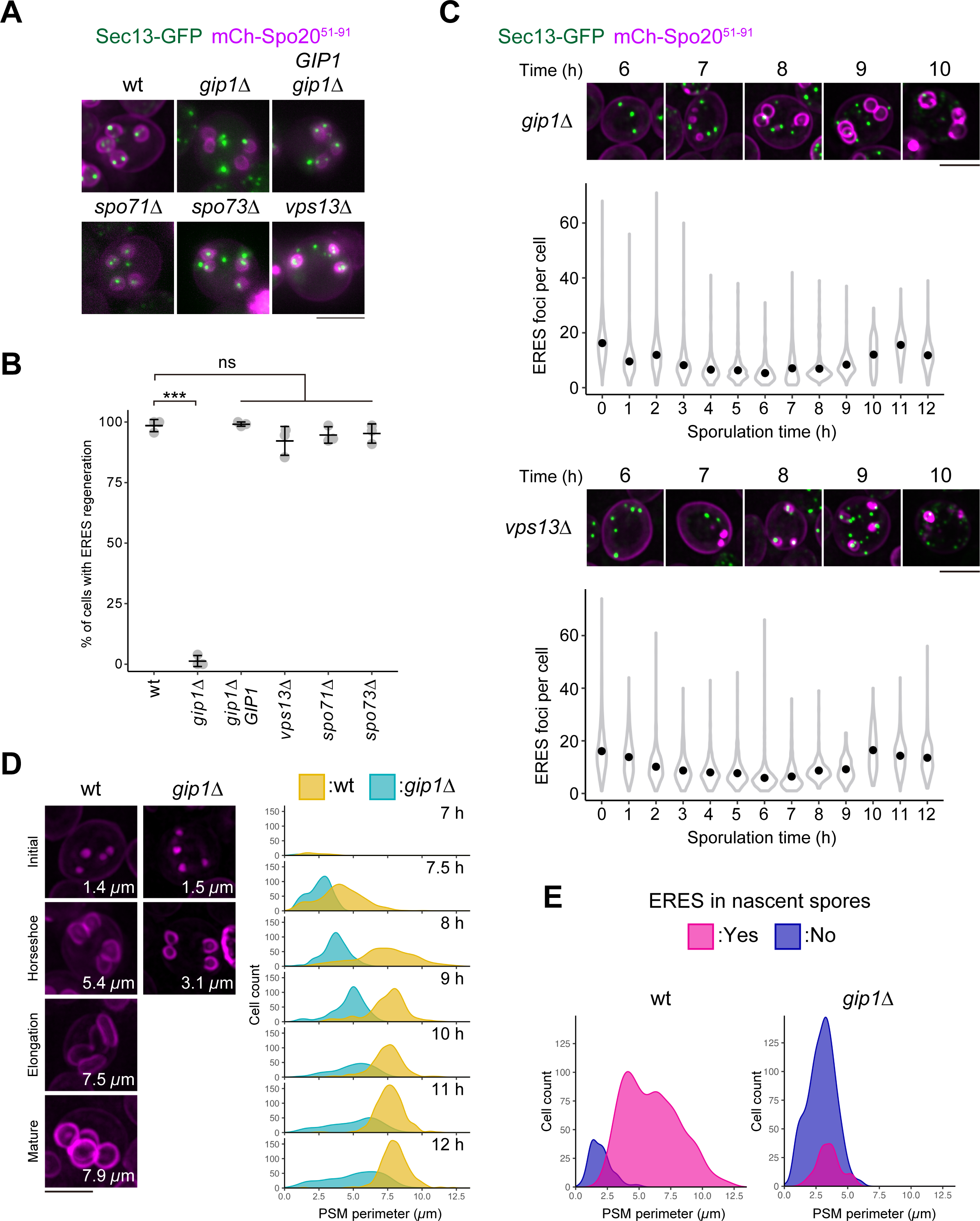
Gip1, a sporulation-specific PP1 targeting subunit, is required for ERES capture by PSMs. (A) Maximum intensity projections of cells with the horseshoe-stage of PSMs expressing Sec13-GFP with mCherry-Spo20^51–91^ in wild-type and mutants defective for PSM growth, *gip1*Δ, *gip1*Δ with *CEN-GIP1*, *spo71*Δ, *spo73*Δ, or *vps13*Δ. (B) Quantification of the experiments in panel A. Graphs show the percentage of cells with PSMs harboring ERES dots. More than 100 cells were analyzed in three independent trials. *P* values were calculated with one-way ANOVA Tukey test. ***, *P* < 0.01, ns, not significant. (C) Maximum intensity projections of mutant cells, *gip1*Δ and *vps13*Δ, expressing Sec13-GFP with mCherry-Spo20^51–91^ at each time point after induction of meiosis. Quantifications of the experiments are shown as number of Sec13-GFP dots at indicated time points (n > 300 cells /h). Data are presented as violin plots with median. (D) Representative maximum intensity projection images of wild-type and *gip1*Δ cells expressing mCherry-Spo20^51–91^ at various stages of PSM growth. Numbers indicate average PSM perimeter. Quantification of the distributions of PSM length per cell at each time point is shown on the right. More than 100 cells were analyzed in each time points except 7 h since only a small number of cells showed PSM signals at this time point. (E) Quantification of the distribution of PSM length and ERES acquisition per cell at 7-8 h after meiosis induction in wild-type and *gip1*Δ (n > 600 cells).

**Figure 5.**
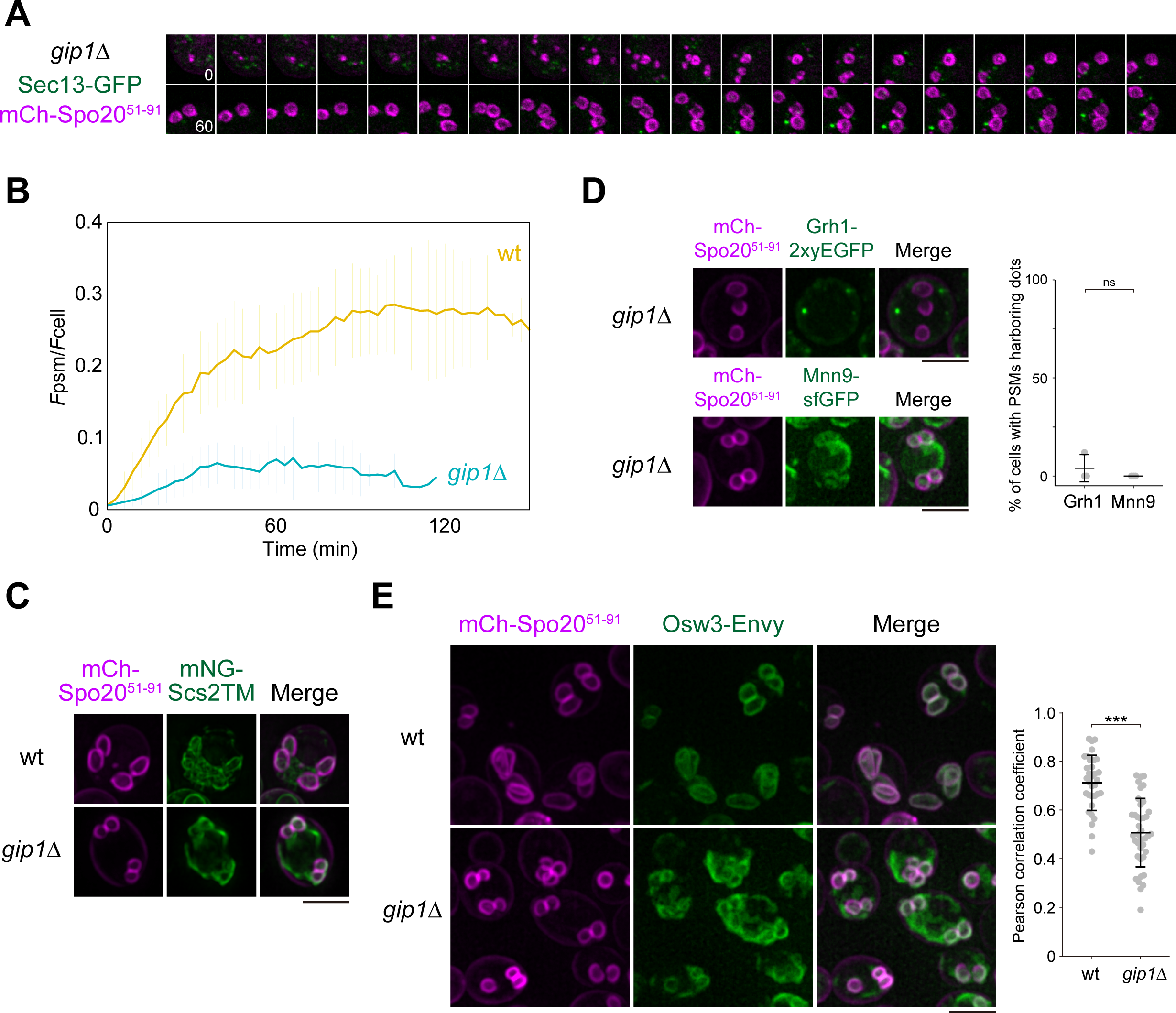
Gip1-dependent regeneration of ERES within presumptive spores. (A) Representative time-lapse image of single presumptive spores from *gip1*Δ cells expressing Sec13-GFP and mCherry-Spo20^51–91^ in meiosis. Kymograph from Movie S2 is shown. (B) Quantification of signal intensities of ERES, Sec13-GFP in the cytoplasm of single PSMs of wild-type and *gip1*Δ cells over time. (C) Maximum intensity projection images of anaphase II cells depicting the ER marked with mNG-Scs2TM and PSM with mCherry-Spo20^51–91^ in wild-type and *gip1*Δ. (D) Maximum intensity projections of the *gip1*Δ cells expressing ERGIC marker Grh1-2xyEGFP and *cis*-Golgi localized enzyme Mnn9-sfGFP with PSM marker mCherry-Spo20^51–91^. Graphs showing the percentage of cells with PSMs harboring indicated markers. More than 100 cells were analyzed in three independent trials. *P* values were calculated with unpaired two-tailed Welch’s *t*-test. (E) Maximum intensity projections of wild-type and the *gip1*Δ cells expressing secretory cargo protein Osw3-Envy and PSM marker mCherry-Spo20^51–91^. Graphs showing the Pearson correlation coefficient values between indicated markers. More than 30 cells were analyzed. *P* values were calculated with unpaired two-tailed Welch’s *t*-test. ***, *P* < 0.01.

To determine whether the observed ERES regeneration defect in *gip1*Δ cells is caused by a failure to properly localize Glc7 or a specific function of Gip1 itself, we next examined the localization of GFP-Glc7 in *gip1*Δ cells, and ERES regeneration in the *glc7-136* mutant, in which the Glc7 protein is defective in association with Gip1^39^. In wild-type cells, GFP-Glc7 was first seen as dots inside the small circular PSM, which may reflect transient SPB localization. Then GFP-Glc7 showed septin-like parallel bars along with the growing PSM, and finally appeared in the nucleus after PSM maturation (Fig. 6A). These localization patterns of GFP-Glc7 are consistent with the dynamic localization of Gip1 during PSM formation, which we have shown previously^25^. In *gip1*Δ cells, SPB-localization of GFP-Glc7 was normal, but the subsequent characteristic localizations along the PSM and septin was not observed (Fig. 6B), confirming our previous observation^26^. Moreover, ERES regeneration defect was similarly observed in the *glc7-136* mutant allele (Fig. 6C). Previously, we mapped the septin-localization domain in Gip1, and the deletion of this domain (Gip1-Δsep) resulted in protein localization at the PSM instead of proper localization with septin^25^, but neither PSM growth nor sporulation defects were seen under this condition. Consistently, Gip1-Δsep was able to completely suppress the ERES regeneration defect in *gip1*Δ cells, as the wild-type version of Gip1 did (Fig. S2B). Thus, the defect in ERES regeneration in *gip1*Δ can be the consequence of the failure to recruit Glc7 to the PSM during PSM formation.

**Figure 6.**
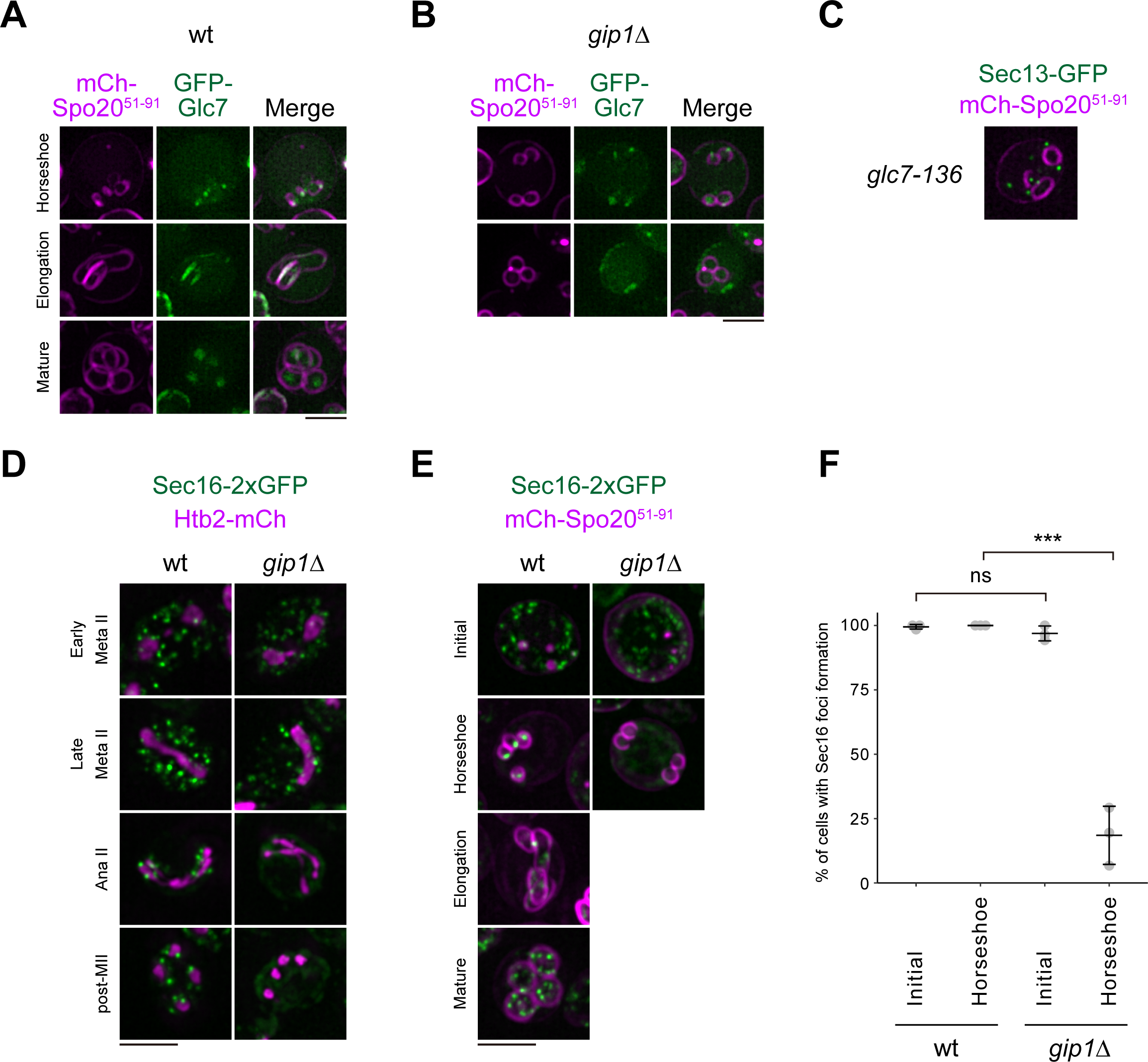
Gip1-dependent foci formation of Sec16 at the onset of PSM formation. (A-B) Maximum intensity projections of cells with various stages of PSMs expressing GFP-Glc7 with mCherry-Spo20^51–91^ in wild-type (A) and *gip1*Δ (B). (C) Representative maximum intensity projection image of Sec13-GFP and mCherry-Spo20^51–91^ in *glc7-136* cells, in which the association is defective between Glc7, the catalytic subunit of PP1, and Gip1, a development-specific targeting subunit of PP1. (D-E) Maximum intensity projections of cells at indicated stages during meiotic progression. Wild-type and *gip1*Δ cells expressing Sec16-2xyEGFP with (D) Htb2-mCherry or (E) mCherry-Spo20^51–91^. (F) Quantification of the experiments shown in panel E. More than 30 cells were analyzed in three independent trials. *P* values were calculated with one-way ANOVA Tukey test. ***, *P* < 0.01, ns, not significant.

The next question is how the PP1-Gip1 signal is transmitted to ERES formation? We hypothesize that the factor organizing ERES is regulated by the PP1-Gip1 during sporulation and Sec16 is a candidate. In wild-type cells, Sec16-2xyEGFP foci were dispersed throughout cytoplasm and distributed along the nucleus at anaphase II, which underwent alterations in meiotic progression, and were precisely segregated into the nascent spore cytoplasm, consistent with the observation of ERES visualized with Sec13-GFP (Fig. 6D, 6E). In *gip1*Δ cells, we found that Sec16-2xyEGFP was unable to form foci upon transition from the initial to horseshoe-stage of PSM formation (Fig. 6F). Thus, whether directly or indirectly, PP1-Gip1 transduces signals toward ERES regeneration via development of Sec16 foci during PSM formation.

### Defect in ERES formation also affects PSM extension

Yorimitsu *et al.* have revealed that Sec16 foci formation is mediated through the interaction with Sed4^40^. Sed4 is a non-essential paralog of Sec12, the ER-resident transmembrane guanine-nucleotide-exchange-factor (GEF) for Sar1 GTPase, which has no catalytic activity toward Sar1, but is proposed to stimulate Sar1 activity^41–43^. We found that a single deletion mutant for *SED4* has partial defect in sporulation (Fig. 7A). Consistent with the proposed mitotic localization of Sec12 and Sed4, at the entire ER and dot-like signals on the ER, respectively^44^, similar localizations of sfGFP-Sec12 and sfGFP-Sed4 were observed during sporulation (Fig. 7B). When we observed the Sec13-GFP and mCh-Spo20^51–91^ in *sed4*Δ, a significant fraction of the *sed4*Δ cells showed defective ERES segregation and unequal, aberrant PSM extension (Fig. 7C, 7D). These defects were suppressed by introducing either full-length Sed4 or Sed4 lacking the luminal domain, which is dispensable for its function^41^ (Fig. 7C). Live-cell imaging of ERES and the PSM by SCLIM showed that, in *sed4*Δ cells, ERES formation was normal within the PSM that showed a relatively normal extension rate, but was defective within PSMs defective in extension (Fig. 7E, 7F, and Movie S3). These results again confirmed that the *de novo* formation of ERES is indeed required for the subsequent extension of PSMs. As shown in Fig. 6F, Sec16 foci formation was defective in *gip1*Δ cells. Likewise, dot-like localization pattern of sfGFP-Sed4 inside PSMs was also defective in *gip1*Δ cells (Fig. 7G). This is consistent with the finding of mutual localization between Sec16 and Sed4^40^. These observations suggest that Sed4 together with Sec16 is also involved in ERES regulation by PP1-Gip1.

**Figure 7.**
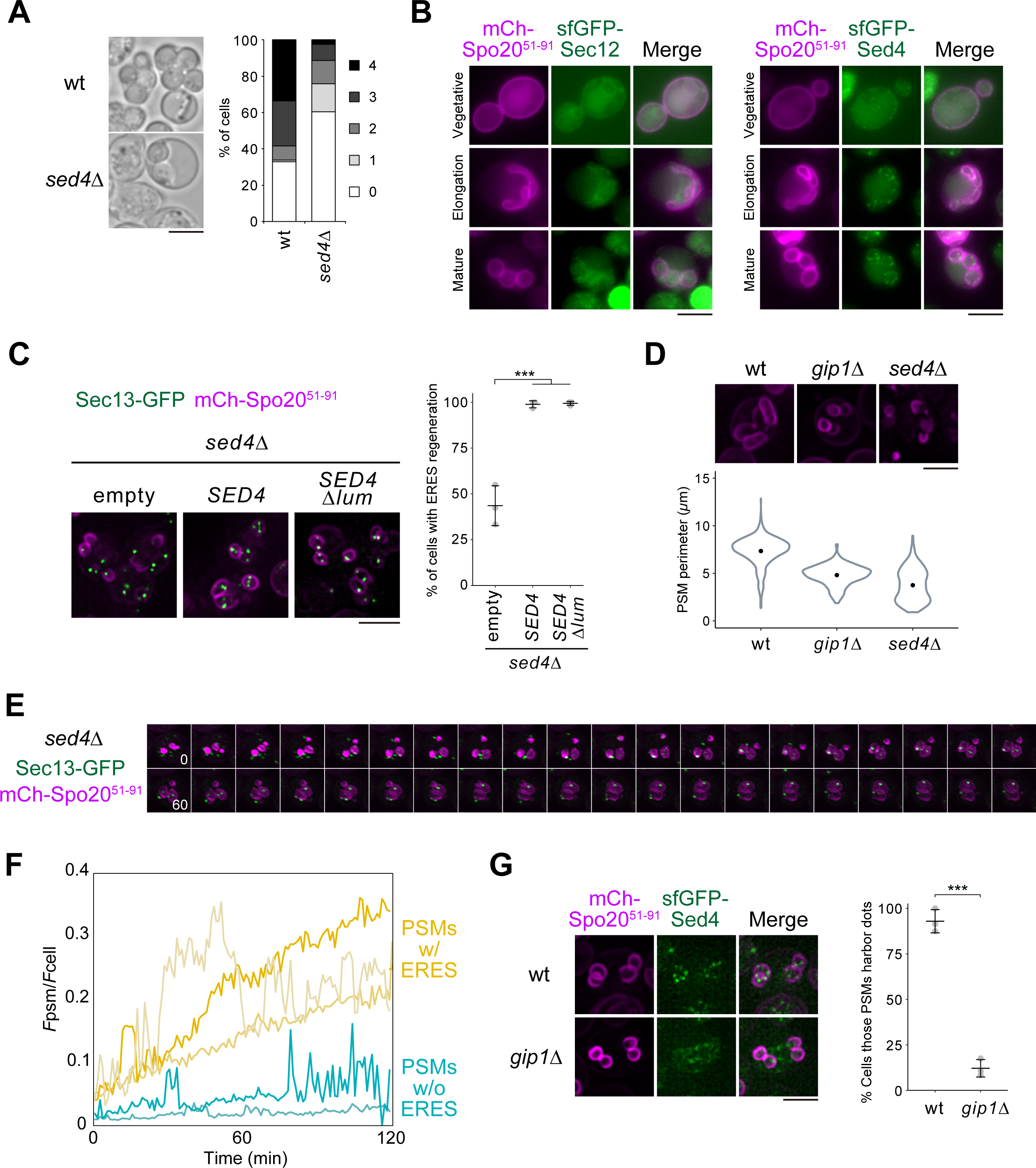
Deletion of *SED4* phenocopied most of the defects observed in *gip1*Δ cells. (A) Representative bright field images of wild-type and *sed4*Δ cells that were induced to sporulate. Graph shows the quantification of the distribution of asci in left panels. (B) Representative images of GFP-Sec12 or GFP-Sed4 with mCherry-Spo20^51–91^ at each stage of meiosis in wild-type cells. (C) Maximum intensity projection images of cells with Sec13-GFP and mCherry-Spo20^51–91^ in *sed4*Δ harboring empty vector, full-length *SED4*, or *SED4* lacking the luminal domain (*SED4*Δ*lum*). Quantification of the experiments shown in left panels is on the right. More than 50 cells were analyzed in three independent trials. *P* values were calculated with one-way ANOVA Tukey test. ***, *P* < 0.01. (D) Maximum intensity projection images of wt, *gip1*Δ, and *sed4*Δ cells with mCherry-Spo20^51–91^. Quantification of the PSM perimeters in wild-type, *gip1*Δ, and *sed4*Δ (n > 600 cells). Data are presented as violin plots with median. (E) Representative time-lapse image of single presumptive spores from *sed4*Δ cells expressing Sec13-GFP and mCherry-Spo20^51–91^ in meiosis. Kymograph from Movie S3 is shown. (F) Quantification of signal intensities of ERES, Sec13-GFP in the cytoplasm of single PSMs of wild-type and *gip1*Δ cells over time. (G) Maximum intensity projection images of cells with GFP-Sed4 and mCherry-Spo20^51–91^ in wt and *gip1*Δ. Graphs showing the percentage of cells with PSMs harboring sfGFP-Sed4 dots. More than 30 cells were analyzed in three independent trials. *P* values were calculated with unpaired two-tailed Welch’s *t*-test.

## Discussion

We investigated the dynamics of organelles along the secretory pathway in budding yeast during the formation of *de novo* membranes in meiosis. In this analysis, we found that the number of ERES, the starting point of membrane traffic, decreased, and then increased after anaphase II. The increase of the ERES number is due to their regeneration, which is restricted within the cytoplasmic space surrounded by the newly formed PSM. Furthermore, we showed that ERES regeneration could be mediated through Sec16 and is triggered by PP1-Gip1, which is specifically targeted to the inner surface of PSMs during meiosis. A mutant defective in both COPII carrier and ERES formation showed the PSM growth defect that is reminiscent of the defect in *gip1*Δ cells. These observations suggest that the involvement of early secretory pathway in the formation of PSMs may be regulated by the phosphorylation. In summary, our results have revealed that developmentally regulated ERES regeneration is essential for PSM growth by mediating segregation of the organelles that constitute membrane traffic, including the Golgi, which presumably supplies membrane lipids to PSMs through the secretory pathway (Fig. 8). This process may also contribute to the renewal of organelles along the secretory pathway that have been damaged through aging, leading to maintenance of proteostasis.

**Figure 8.**
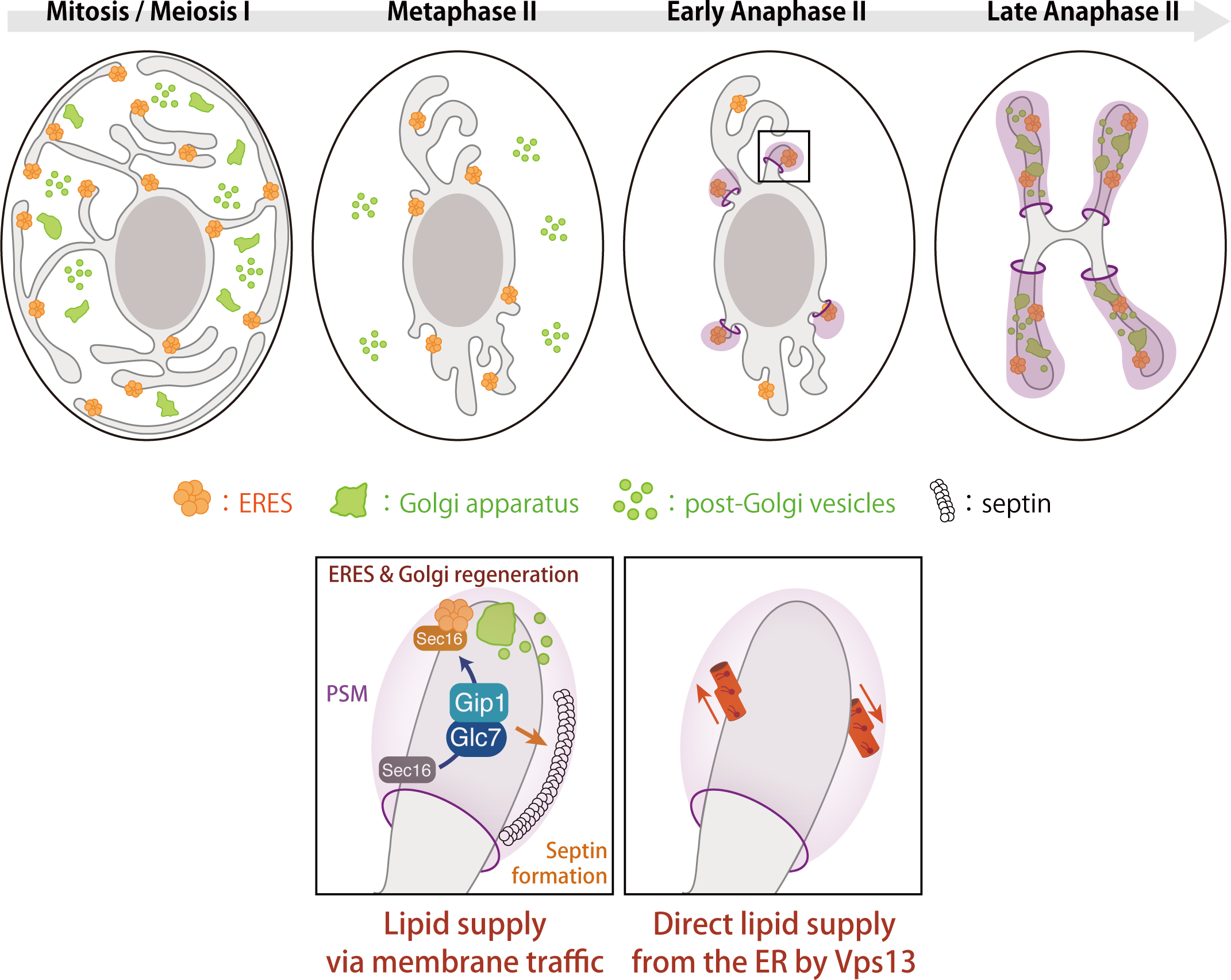
Model for the early step of PSM formation. Coalescence and fusion of post-Golgi vesicles initiate nascent PSM formation at SPBs. ERES formation could be transiently inactivated during meiosis, then ERES regeneration is locally activated in the region surrounded by the PSM in early anaphase II. Gip1-Glc7 mediates ERES regeneration presumably through the foci-formation of Sec16, Golgi re-assembly, and supplies membrane lipids toward PSMs via membrane traffic. Gip1-Glc7 is also essential for septin high-order structural formation in the spore cytoplasm. In the process of PSM extension, the Vps13-complex also functions as the direct lipid conduit from the ER to PSMs.

### Meiosis-specific ERES regeneration leads to both PSM formation and segregation of the secretory organelles into gametes

In mitosis, the number of ERES fluctuates, according to the processes of assembly and disassembly, and fusion and fission^13–15,45^. In the present study, we have shown that the ERES numbers gradually decrease as meiosis progresses, and then increase after the onset of PSM formation due to local regeneration in the nascent spore cytoplasm. The ERES dynamics and anomaly of local ERES regeneration in *gip1*Δ mutant cells led us to propose that there is a signal that directly or indirectly represses ERES formation in the early stages of meiosis. At the onset of PSM formation, however, ERES formation is de-repressed in a Gip1-dependent manner. The key molecule is Sec16, because Sec16 is the most upstream and the important regulator of ERES formation^14,15^. Although numerous phosphorylation sites have been found in Sec16, their specific roles remain unclear^46^. In mammalian cells, secretion mediated by membrane traffic pauses during mitosis^47^. This phenomenon is due to the disruption of the early secretory pathway that includes ERES and the Golgi, and some parts of the mechanism have been proposed recently^18^. ERES formation in mammals is determined by the interaction between Sec16 and TANGO1, and the affinity of TANGO1 for Sec16 is regulated by phosphorylation in a cell cycle-dependent manner. The phosphorylation status of TANGO1 is in equilibrium through the action of CK1 and PP1, but PP1 activity is specifically decreased during mitosis, leading to the dissociation of Sec16 from the ER and ERES disassembly^17^. Although TANGO1 is absent in budding yeast, development-specific regulation of Sec16 recruitment could exist. We found that the mutant of Sed4, a regulator of the Sar1 GTPase cycle and Sec16 localization to ERES^40,41^, partially phenocopied *gip1*Δ, suggesting involvement of these factors in the ERES formation during meiosis.

Our previous analysis of organellar segregation in sporulation is inconclusive as to whether the Golgi segregation into spores is mediated through the active transport or PSMs act as a diffusion barrier for the Golgi^20^. In the present study, we showed that local regeneration of ERES and the Golgi occurs specifically in the spore cytoplasm in wild-type cells, but neither was observed in *gip1*Δ cells. Thus, the Golgi apparatus is once dissipated from the cell and reassembled specifically within the nascent spores during meiosis, and this phenomenon appears equivalent to the mitotic disassembly and reassembly of the Golgi in mammalian cells. How the Golgi apparatus is formed and maintained in cells remains elusive. To clarify such questions, live cell observations of the Golgi regeneration were conducted in yeast and plant cells using a drug that induces Golgi disassembly followed by its washout^30,48,49^. Results from these studies showed that the Golgi enzymes are absorbed into the ER upon disassembly and return to the Golgi through ERES when the Golgi is reassembled by drug washout. Moreover, in plant cells, a subset of the Golgi-resident proteins underwent nucleation near ERES as a template for Golgi regeneration after drug removal. The authors concluded that it served as the scaffold for Golgi regeneration and named it Golgi entry core compartment (GECCO)^48,50^. GECCO is functionally equivalent to ERGIC of mammals and their presence has been demonstrated in yeast^30^. Considering the evolutional conservation of ERGIC/GECCO as an earliest compartment for Golgi regeneration, they may be segregated first into the spore cytoplasm during PSM formation. The molecular mechanism underlying this phenomenon in meiosis is clearly regulated by PP1-Gip1, and similar mechanisms might be involved in Golgi regeneration in mitosis.

PP1-Gip1 is also essential for the formation of the septin higher-order structure beneath PSMs^26^. We showed that the Gip1 mutant that lacks previously identified septin localization domain (Gip1Δsep) can also trigger ERES formation and PSM formation^25^. This result is in agreement with our previous report, in which the localization of PP1-Gip1 around PSMs is sufficient for its function^25^. The observed sporulation defects in *gip1*Δ cells can be explained by Gip1’s function as a trigger for membrane traffic remodeling and its contribution to PSM formation.

### Nutrient starvation-triggered meiotic remodeling in yeast

In flies and mammals, nutrient starvation is shown to result in the formation of a membrane-less structures of Sec16, called Sec bodies, by liquid-liquid phase separation together with a subset of COPII components^19,51^. Sec bodies are proposed to act as temporal reservoirs for COPII components when secretion is blocked in response to stress, that is immediately resolved upon stress removal^51^. Although yeast meiosis is a process triggered by starvation, our results have shown that ERES are reduced in number but do not undergo the formation of membrane-less structures; ERES remain active during meiosis.

King et al. found that the mechanism of cell rejuvenation following sporulation is achieved through the elimination of senescence factors such as non-chromosomal rDNA circles with nuclear pore complexes from the gametes, and that PSMs are key for this elimination^24^. In the present study, we found PP1-Gip1-triggered reorganization of membrane traffic specifically in the cytoplasmic space of PSMs, and cells may use this mechanism to eliminate senescence factors.

Our study provides a framework for the reorganization of membrane traffic during meiosis to form PSMs. This elaborate maneuver not only generates *de novo* membrane structures but also partitions the ERES, the Golgi, and ultimately the entire membrane traffic into new cytosolic compartments created by the PSM. Importantly, a comparison of lipid supply to the PSM with autophagosome and acrosome membrane formation highlights the facts that these membrane structures are generally cup-shaped bilayer structures with a lumen, derived from vesicles from the Golgi, and their growth is directly mediated by lipid transporters including Vps13 and Atg2^52,53^. During the formation of the autophagosomes, the proximity of the autophagosomal isolation membrane to the ERES has been observed, and direct fusion of COPII vesicles has also been proposed as a source of membrane lipids^54,55^. Thus, the cell can utilize various strategies to assemble nascent membranes from scratch. Identifying these individual mechanisms and uncovering their shared features will provide key insights into both areas.

## Materials and methods

### Yeast strains and plasmids

Standard genetic techniques were used unless otherwise noted. All strains used in this study are derivatives of SK1. Strains and plasmids are listed in Table S1 and S2, respectively. The following alleles and plasmids were constructed in previous studies: *NDT80::hphNT1::P_4xlexA_-9xMyc-NDT80* and *AUR1::P_ACT1_-LexA-ER-haVP16::AUR1-C*^5^, *mTagBFP2-Spo20*^51–915^, and *glc7-136*^26^.

Deletion and tagging of genes at their endogenous loci were performed by standard PCR-based methods^55–58^. The pNC160-GFP-*GLC7* was a gift from K. Tatchell^59^.

To construct *pRS306-mCherry-Spo20*^51–91^ as a marker for PSM, sequences for mCherry and a fragment of Spo20 were cloned into a *URA3* integrating vector harboring the *TEF1* promoter and *DIT1* terminator. For the visualization of nucleus, *Htb2-mCherry*^8^ was cloned into *URA3* integrating vector. To generate *pRS303-mNG-Scs2TM*, sequences for mNeonGreen and a transmembrane domain of Scs2 were cloned into the *HIS3* integrating vector harboring the *TEF1* promoter and *DIT1* terminator. The full-length gene including upstream and downstream flanking regions encoding *GIP1* was cloned into the *HIS3 CEN* vector. To construct *HIS3* integration plasmids harboring either full-length *GIP1* or *GIP1*Δ*sep*, corresponding sequences from *CEN TRP1* plasmids^25^ were digested and cloned into pRS303. All *HIS3* integration plasmids were integrated into the genome by BssHII digestion. Similarly, *URA3* integration constructs were integrated by cutting with EcoRV. Coding sequences for Sed4 or Sed4 without the luminal domain were cloned into the *HIS3* integration vector together with upstream and downstream flanking regions, and transformed into *sed4*Δ*::kanMX6* strains.

### Growth conditions

Unless otherwise noted, standard media were used^60^. Induction of sporulation was performed essentially as described^4^. Synchronous sporulation induction by the estradiol-inducible *NDT80* block/release system was carried out as described previously^5,61^.

### Fluorescence microscopy and image analyses

Static images were acquired with a Leica Thunder Imager Live Cell equipped with a Leica HC APO 100x oil immersion lens (NA 1.4), sCMOS camera DFC9000 GTC, as well as by using a Keyence BZ-X710. Time-lapse imaging was performed with a super-resolution confocal live imaging microscopy (SCLIM2), which we developed based on our basic imaging system SCLIM^27,28^. This system consists of a Nikon ECLIPSE Ti2-E inverted microscope equipped with a NIKON APO TIRF 100x NA 1.49 oil immersion objective lens, a high-speed spinning-disk confocal scanner (CSU-X1, Yokogawa Electric), a custom-made spectroscopic unit, an image intensifier (Hamamatsu Photonics) with a custom-made cooling system, and three sCMOS cameras (Zyla 5.5, Andor) for green, red, and infrared observation. Image acquisition was executed by NIS-Elements (Nikon). For 3D (*xyz*) and 4D (*xyz* plus time) observations, cells induced for sporulation were immobilized on a glass-based dish (Iwaki) by sandwiching with a 1% agarose pad containing 1% KOAc and imaged. Stacks of ∼ 50 image planes were collected with a spacing of 0.2 µm to cover the entire cell. Z-stack images were subjected to deconvolution (iterative restoration) with Volocity software (Perkin Elmer) using the theoretical point-spread function for spinning-disk confocal microscopy, then combined using maximum intensity z-projection.

### Image quantification

Intensity profile images for Sec13-GFP were obtained using the Metamorph built-in module (Molecular Devices). For the quantification analysis of ERES numbers, Sec13-GFP images were segmented into background and objects using the ilastik pixel classification workflow^62^. Cell masks were also generated from bright field images using CellPose^63^. Distribution of the numbers for segmented ERES pixels within the area of the cell was analyzed by Fiji and custom script in R. PSM perimeter was analyzed by Fiji. For time-course analysis of ERES intensities, the PSM of interest was tracked manually and fluorescent intensity (*F_PSM_*) for the green channel was averaged within the region for PSM and normalized to averaged fluorescent intensity of the whole cell (*F_cell_*) over time.

### FRAP analysis

FRAP analysis was performed using confocal laser scanning microscopy (LSM880, Carl Zeiss) with a 63x Apochromat objective lens (NA 1.4) and a 488-nm laser. The bleaching routine started with 2 pre-bleach scans followed by a bleaching scan, and the recovery of fluorescence was monitored every 1 s for 60 s at 0.2% laser intensity. The fluorescence recovery data were normalized using data acquired from the non-bleached regions. Data were fitted to a double exponential curve to calculate half-time to recovery after full scale normalization using easyFRAP software^64^.

## Acknowledgement

We are grateful to Aaron Neiman for fruitful discussion on this manuscript and all the encouragement through the project. We thank all the members of Molecular Biology Laboratory at Tsukuba University and Live Cell Super-Resolution Imaging Research Team at RIKEN Center for Advanced Photonics for discussions. We also thank Kalai Madhi Muniandy and Miho Waga for technical assistance. This work was supported by Grants-in-Aid for Scientific Research from the Ministry of Education, Culture, Sports, Science, and Technology (MEXT) of Japan (grant numbers 21K06145 to Y.S., 20K05782 and 23K05006 to H.T., 18H05275 to A.N., 22K06074 to K.I.).

## Author contributions

Conceptualization: Y.S.

Formal analysis: Y.S. and T.S.

Investigation: Y.S. and T.S.

Resource: H.T.

Writing-Original Draft: Y.S.

Writing-Review & Editing: Y.S., H.T., K.K., A.N., and K.I.

Visualization: Y.S. and T.S.

---

**Movie S1**

Dynamics of Sec13-GFP and mCherry-Spo20^51–91^ in wild-type cell, related to Figure 1E.

**Movie S2**

Dynamics of Sec13-GFP and mCherry-Spo20^51–91^ in *gip1*Δ cell, related to Figure 5A.

**Movie S3**

Dynamics of Sec13-GFP and mCherry-Spo20^51–91^ in *sed4*Δ cell, related to Figure 7E.

**Figure S1.**
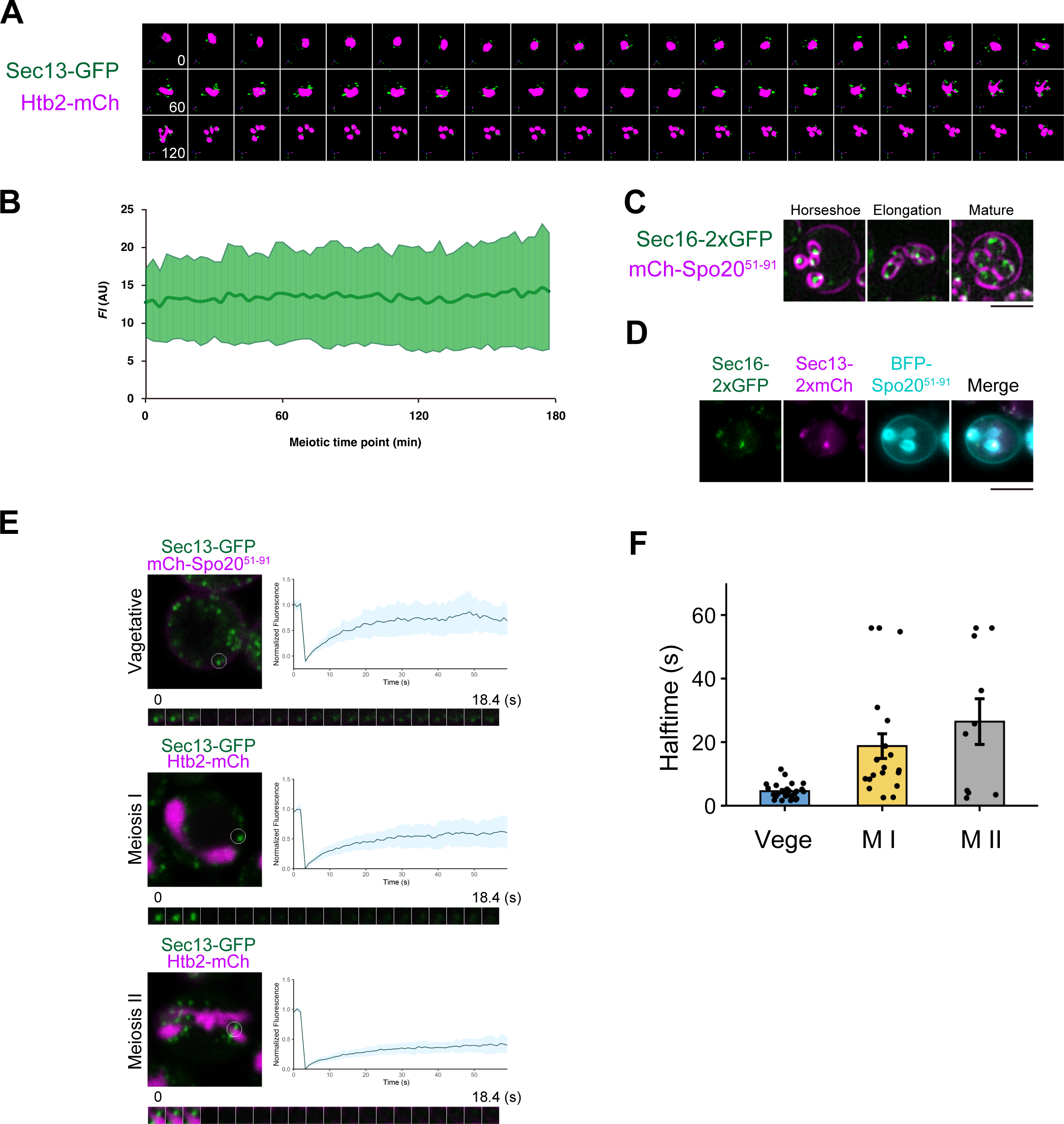
(A) Montage of cells expressing Sec13-GFP and Htb2-mCherry in meiotic progression. Quantification of Sec13-GFP signal intensity over time is also shown. Median is shown as a bold line. (B) Quantification of signal intensity of Sec13-GFP per cell over time. (C) Maximum intensity projections of the cells expressing Sec16-2xyEGFP, mCherry-Spo20^51–91^ in meiosis. (D) Maximum intensity projections of the cells expressing Sec16-2xyEGFP, Sec13-2xmCherry, and tagBFP-Spo20^51–91^ in meiosis. (E) FRAP analysis of Sec13-GFP at ERES in vegetatively growing or sporulating cells. Bleached area (circle) is shown in single plane images of the upper left. Data represents in the upper right, mean ± SEM, n≥10. Montage of Sec13-GFP recoveries is shown in the bottom panels. (F) Quantification of half-time recoveries of the cells in vegetative growth or in each stage of meiotic progression.

**Figure S2.**
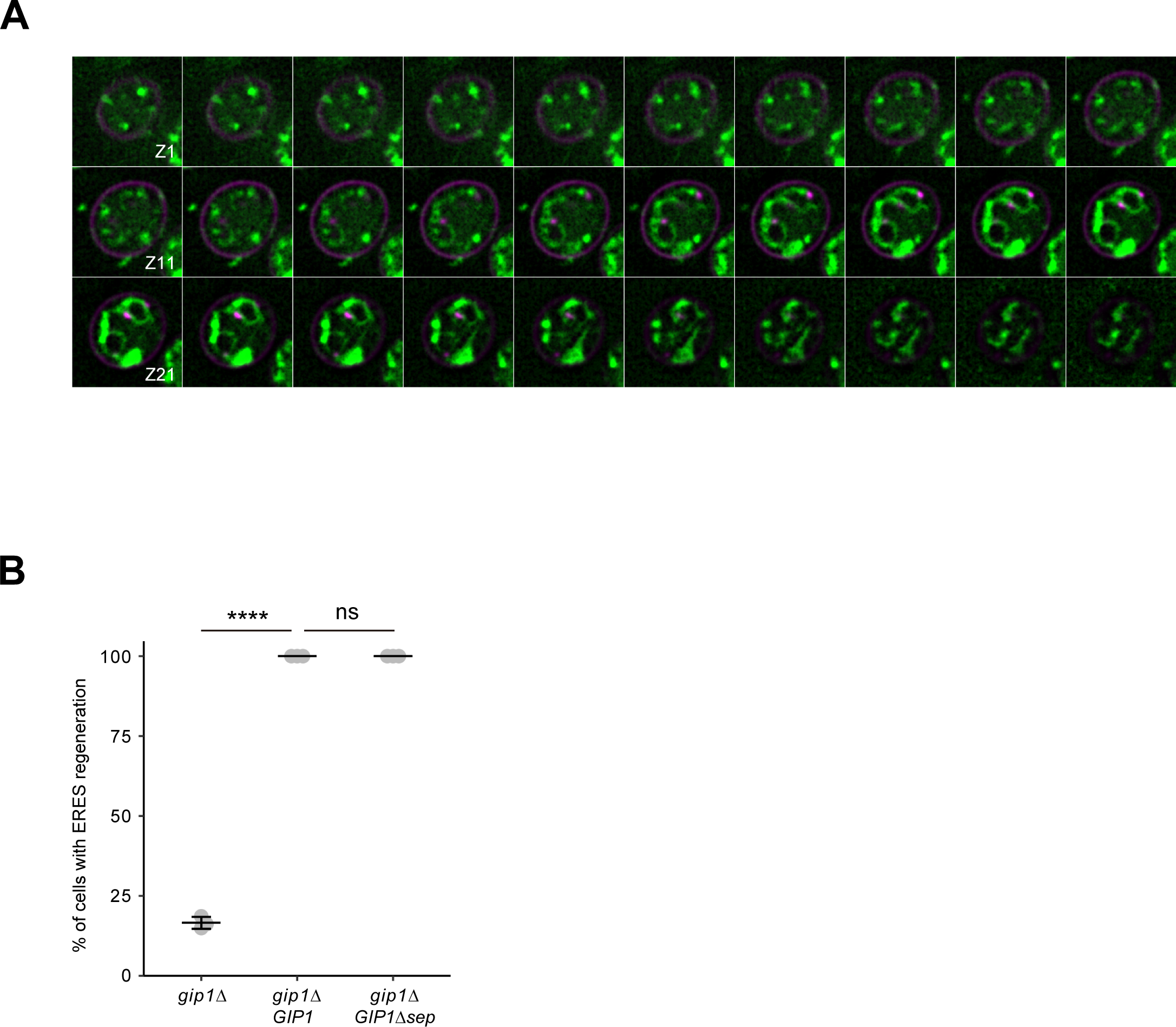
Gip1 contributes to ERES regeneration localizing only in the vicinity of PSMs. Quantification of ERES regeneration in presumptive spores in *gip1*Δ cells expressing full-length Gip1 or Gip1Δsep lacking the septin localization domain.

