## Supplementary material for "Remodeling of the secretory pathway is coordinated with *de novo* membrane formation in budding yeast gametogenesis": Table S1

**S1 Table. Yeast strains used in this study.**

TNY168 was used as background strain unless otherwise noted.

Name	Genotype	Source

TNY168	*MAT***a**/*MAT*α *his3*Δ*SK*/*his3*Δ*SK ura3*/*ura3 trp1::hisG*/*trp1::hisG leu2*/*leu2 arg4-NspI/ARG4 lys2/lys2 ho*Δ*::LYS2/ho*Δ*::LYS2*	^1^

*rme1::LEU2/RME1 AUR1::P_ACT1_-LexA-ER-haVP16::AUR1-C/AUR1::P_ACT1_-LexA-ER-haVP16::AUR1-C*

*NDT80::hphNT1::P4×lexA-9×Myc-NDT80* /*NDT80::hphNT1::P4×lexA-9×Myc-NDT80*

YSIY420	*ura3*/*ura3::pRS306-HTB2-mCherry his3*Δ*SK*/*his3*Δ*SK::pRS303-mNG-SCS2TM*	This study

YSIY961	*SEC13-yEGFP::TRP1/SEC13-yEGFP::TRP1 ura3*/*ura3::pRS306-mCherry-Spo20^51-91^*	This study

YSIY962	*SEC13-yEGFP::TRP1/SEC13-yEGFP::TRP1 ura3*/*ura3::pRS306-HTB2-mCherry*	This study

YSIY835	*SEC16-2xyEGFP::kanMX6/SEC16-2xyEGFP::kanMX6 ura3*/*ura3::pRS306-mCherry-Spo20^51-91^*	This study

YSIY1279	*SEC16-2xyEGFP::kanMX6/SEC16-2xyEGFP::kanMX6 SEC13-2xmCherry::TRP1/SEC13-2xmCherry::TRP1*	This study

YSIY1291	*GRH1-2xyEGFP::kanMX6/ GRH1-2xyEGFP::kanMX6 ura3*/*ura3::pRS306-mCherry-Spo20^51-91^*	This study

YSIY1208	*MNN9-sfGFP::natNT2/MNN9-sfGFP::natNT2 ura3*/*ura3::pRS306-mCherry-Spo20^51-91^*	This study

YSIY1209	*gip1*∆*::kanMX6/gip1*∆*::kanMX6 MNN9-sfGFP::natNT2/MNN9-sfGFP::natNT2 ura3*/*ura3::pRS306-mCherry-Spo20^51-91^*	This study

YSIY1657	*gip1*∆*:: natNT2/gip1*∆*::natNT2 GRH1-2xyEGFP::kanMX6/ GRH1-2xyEGFP::kanMX6 ura3*/*ura3::pRS306-mCherry-Spo20^51-91^*	This study

YSIY820	*rtn1*∆*::kanMX6/rtn1*∆*::kanMX6 rtn2*∆*::kanMX6/rtn2*∆*::kanMX6 yop1*∆*::natNT2/yop1*∆*::natNT2*	This study

*AUR1/AUR1::P_ACT1_-LexA-ER-haVP16::AUR1-C ura3*/*ura3::pRS306-mCherry-Spo20^51-91^*

YSIY821	*rtn1*∆*::kanMX6/rtn1*∆*::kanMX6 rtn2*∆*::kanMX6/rtn2*∆*::kanMX6 yop1*∆*::natNT2/yop1*∆*::natNT2*	This study

*AUR1/AUR1::P_ACT1_-LexA-ER-haVP16::AUR1-C ura3*/*ura3::pRS306-HTB2-mCherry*

YSIY561	*rtn1*∆*::kanMX6/rtn1*∆*::kanMX6 rtn2*∆*::kanMX6/rtn2*∆*::kanMX6 yop1*∆*::natNT2/yop1*∆*::natNT2*	This study

*AUR1/AUR1::P_ACT1_-LexA-ER-haVP16::AUR1-C SEC13/SEC13-2xyEGFP::TRP ura3*/*ura3::pRS306-mCherry-Spo20^51-91^*

YSIY566	*gip1*∆*::kanMX6/gip1*∆*::kanMX6 SEC13-yEGFP::TRP1/SEC13-yEGFP::TRP1 ura3*/*ura3::pRS306-mCherry-Spo20^51-91^*	This study

YSIY592	*spo71*∆*::kanMX6/spo71*∆*::kanMX6 SEC13-yEGFP::TRP1/SEC13-yEGFP::TRP1 ura3*/*ura3::pRS306-mCherry-Spo20^51-91^*	This study

YSIY594	*spo73*∆*::kanMX6/spo73*∆*::kanMX6 SEC13-yEGFP::TRP1/SEC13-yEGFP::TRP1 ura3*/*ura3::pRS306-mCherry-Spo20^51-91^*	This study

YSIY596	*vps13*∆*::kanMX6/vps13*∆*::kanMX6 SEC13-yEGFP::TRP1/SEC13-yEGFP::TRP1 ura3*/*ura3::pRS306-mCherry-Spo20^51-91^*	This study

YSIY556	*ura3*/*ura3::pRS306-mCherry-Spo20^51-91^ his3*Δ*SK::pRS303-mNG-SCS2TM*/*his3*Δ*SK::pRS303-mNG-SCS2TM*	This study

YSIY1157	*gip1*∆*::kanMX6/gip1*∆*::kanMX6 ura3*/*ura3::pRS306-mCherry-Spo20^51-91^ his3*Δ*SK*/*his3*Δ*SK::pRS303-mNG-SCS2TM*	This study

YSIY1680	*OSW3/OSW3-ENVY::natNT2 ura3*/*ura3::pRS306-mCherry-Spo20^51-91^*	This study

YSIY1681	*gip1*∆*::kanMX6/gip1*∆*::kanMX6 OSW3/OSW3-ENVY::natNT2 ura3*/*ura3::pRS306-mCherry-Spo20^51-91^*	This study

YSIY1218	*ura3*/*ura3::pRS306-mCherry-Spo20^51-91^*	This study

YSIY793	*gip1*∆*::kanMX6/gip1*∆*::kanMX6 ura3*/*ura3::pRS306-mCherry-Spo20^51-91^*	This study

YSIY1158	*glc7::HIS3MX6/glc7::HIS3MX6 SEC13/SEC13-2xyEGFP::kanMX6 ura3::pRS306-glc7-136*/*ura3::pRS306-glc7-136*	This study

YSIY1241	*gip1*∆*::kanMX6/gip1*∆*::kanMX6 SEC13-yEGFP::TRP1/SEC13-yEGFP::TRP1 ura3*/*ura3::pRS306-mCherry-Spo20^51-91^*	This study

*his3*Δ*SK::pRS303-GIP1*/*his3*Δ*SK::pRS303-GIP1*

YSIY1242	*gip1*∆*::kanMX6/gip1*∆*::kanMX6 SEC13-yEGFP::TRP1/SEC13-yEGFP::TRP1 ura3*/*ura3::pRS306-mCherry-Spo20^51-91^*	This study

*his3*Δ*SK::pRS303-GIP1∆sep*/*his3*Δ*SK::pRS303-GIP1∆sep*

YSIY350	*SEC13-yEGFP::TRP1/SEC13-yEGFP::TRP1*	This study

YSIY351	*sed4*∆*::kanMX6/sed4*∆*::kanMX6 SEC13-yEGFP::TRP1/SEC13-yEGFP::TRP1*	This study

YSIY576	*sfGFP-SEC12/sfGFP-SEC12 ura3*/*ura3::pRS306-mCherry-Spo20^51-91^*	This study

YSIY1354	*sfGFP-SED4/sfGFP-SED4 ura3*/*ura3::pRS306-mCherry-Spo20^51-91^*	This study

YSIY1355	*gip1*∆*::natNT2/gip1*∆*::natNT2 sfGFP-SED4/sfGFP-SED4 ura3*/*ura3::pRS306-mCherry-Spo20^51-91^*	This study

YSIY597	*sed4*∆*::kanMX6/sed4*∆*::kanMX6 SEC13-yEGFP::TRP1/SEC13-yEGFP::TRP1 ura3*/*ura3::pRS306-mCherry-Spo20^51-91^*	This study

YSIY599	*sed4*∆*::kanMX6/sed4*∆*::kanMX6 SEC13-yEGFP::TRP1/SEC13-yEGFP::TRP1 ura3*/*ura3::pRS306-mCherry-Spo20^51-91^*	This study

*his3*Δ*SK::pRS303-SED4*/*his3*Δ*SK::pRS303-SED4*

YSIY601	*sed4*∆*::kanMX6/sed4*∆*::kanMX6 SEC13-yEGFP::TRP1/SEC13-yEGFP::TRP1 ura3*/*ura3::pRS306-mCherry-Spo20^51-91^*	This study

*his3*Δ*SK::pRS303-SED4∆lum*/*his3*Δ*SK::pRS303-SED4∆lum*

YSIY1289	*gip1*∆*::natNT2/gip1*∆*::natNT2 SEC16-2xyEGFP::kanMX6/SEC16-2xyEGFP::kanMX6 ura3*/*ura3::pRS306-HTB2-mCherry*	This study

YSIY1290	*gip1*∆*::natNT2/gip1*∆*::natNT2 SEC16-2xyEGFP::kanMX6/SEC16-2xyEGFP::kanMX6 ura3*/*ura3:: pRS306-mCherry-Spo20^51-91^*	This study

ANS16-1B	*MAT*α *ura3 trp1 his leu2 sec16-2*	^2^

YSIY1664	ANS16-1B, *GRH1-2xyEGFP::kanMX6*	This study

YSIY1665	ANS16-1B, *MNN9-sfGFP::natNT2*	This study

1.	Nakamura, T.S., Suda, Y., Muneshige, K., Fujieda, Y., Okumura, Y., Inoue, I., Tanaka, T., Takahashi, T., Nakanishi, H., Gao, X.-D., et al. (2021). Suppression of Vps13 adaptor protein mutants reveals a central role for PI4P in regulating prospore membrane extension. PLoS Genet. *17*, e1009727. 10.1371/journal.pgen.1009727.

2.	Sato, M., Sato, K., and Nakano, A. (2002). Evidence for the intimate relationship between vesicle budding from the ER and the unfolded protein response. Biochem. Biophys. Res. Commun. *296*, 560–567. 10.1016/s0006-291x(02)00922-1.
