## Supplementary material for "Remodeling of the secretory pathway is coordinated with *de novo* membrane formation in budding yeast gametogenesis": Table S2

**S2 Table. Plasmids used in this study.**

Name	Description	Source

pRS306-mCherry-Spo20^51-91^	*URA3, integration, P_TEF1_-mCherry-Spo20^51-91^*	This study

pRS306-Htb2-mCherry	*URA3, integration, Htb2-mCherry*	This study

pRS303-mNG-Scs2TM	*HIS3, integration, P_TEF1_-mNeonGreen-Scs2^223-244^*	This study

pRS313-BFP-Spo20^51-91^	*HIS3, CEN-ARS, P_TEF1_-mTagBFP2-Spo20^51-91^*	^1^

pRS313-GIP1	*HIS3, CEN-ARS, GIP1*	This study

pRS303-GIP1	*HIS3, integration, GIP1*	This study

pRS303-GIP1∆sep	*HIS3, integration, GIP1∆sep*	This study

pRS303-SED4	*HIS3, integration, SED4*	This study

pRS303-SED4∆lum	*HIS3, integration, SED4∆lim*	This study

pNC160-GFP-GLC7	*TRP1, CEN-ARS, GFP-Glc7*	^2^

1.	Nakamura, T.S., Suda, Y., Muneshige, K., Fujieda, Y., Okumura, Y., Inoue, I., Tanaka, T., Takahashi, T., Nakanishi, H., Gao, X.-D., et al. (2021). Suppression of Vps13 adaptor protein mutants reveals a central role for PI4P in regulating prospore membrane extension. PLoS Genet. *17*, e1009727. 10.1371/journal.pgen.1009727.

2.	Bloecher, A., and Tatchell, K. (2000). Dynamic localization of protein phosphatase type 1 in the mitotic cell cycle of Saccharomyces cerevisiae. J. Cell Biol. *149*, 125–140. 10.1083/jcb.149.1.125.
